# CX3CL1 action on microglia protects from diet-induced obesity by restoring POMC neuronal excitability and melanocortin system activity impaired by high-fat diet feeding

**DOI:** 10.1101/2021.11.08.467521

**Authors:** Jineta Banerjee, Mauricio D. Dorfman, Rachael Fasnacht, John D. Douglass, Alice C. Wyse-Jackson, Andres Barria, Joshua P. Thaler

## Abstract

**Objective:** Diet-induced obesity (DIO) is associated with hypothalamic microglial activation and dysfunction of the melanocortin pathway, but the molecular mechanisms linking the two remain unclear. Previous studies have hypothesized that microglial inflammatory signaling is linked with impaired pro-opiomelanocortin (POMC) neuron function, but this mechanism has never been directly tested *in vivo*. We addressed this hypothesis using the specific microglial silencing molecule, CX3CL1 (fractalkine), to determine whether reducing hypothalamic microglial activation can restore POMC/melanocortin signaling in the brain to protect against DIO.

**Methods:** We performed metabolic analyses in mice with targeted viral overexpression of CX3CL1 in the hypothalamus exposed to high fat diet (HFD). Electrophysiologic recording in hypothalamic slices from POMC-MAPT-GFP mice was used to determine the effects of HFD feeding and microglial silencing via minocycline or CX3CL1 on GFP-labeled POMC neurons. Finally, mice with hypothalamic overexpression of CX3CL1 received central treatment with the melanocortin receptor antagonist SHU-9119 to determine whether melanocortin signaling is required for the metabolic benefits of CX3CL1.

**Results:** We found that targeted expression of both soluble and membrane-bound forms of CX3CL1 in the mediobasal hypothalamus potently reduced weight gain and increased leptin sensitivity in animals exposed to high fat diet. The protective effect of CX3CL1 rescued diet-induced changes in POMC neuron excitability and required intact melanocortin receptor signaling *in vivo*.

**Conclusion:** Our results provide the first evidence that HFD-induced POMC neuron dysfunction involves microglial activation. Furthermore, our study suggests that the anti-obesity action of CX3CL1 is mediated through the restoration of POMC neuron excitability and melanocortin signaling.

## 1. Introduction

Obesity is a leading cause of morbidity and mortality worldwide with few effective treatments [1]. Despite significant research efforts, the mechanisms underlying obesity pathogenesis remain unclear. Current evidence suggests that diet-induced obesity (DIO) is associated with dysfunction of pro-opiomelanocortin (POMC) neurons, a hypothalamic cell population that maintains energy homeostasis by modulating food intake and energy expenditure [2–4]. POMC neurons are activated by nutrients and hormones such as the adipokine leptin, reducing food intake through synaptic release of neurotransmitters (mostly glutamate) and alpha-melanocyte stimulating hormone (α-MSH), a neuropeptide melanocortin 3/4 receptor (MC3/4R) agonist [5–7]. Genetic defects in the leptin-melanocortin pathway such as mutation in MC4R lead to profound high-fat diet (HFD) sensitivity and increased obesity susceptibility in rodents and humans [2,8–12]. Likewise, nongenetic POMC neuron defects acquired during HFD exposure involving α-MSH production, synaptic reorganization, and intracellular calcium responses [3,4] likely contribute to DIO pathogenesis [3,13,14].

In addition to changes in neuronal activity, obesity pathogenesis involves chronic hypothalamic inflammation as a critical component [15–21]. Activation of microglia (CNS macrophage-like immune cells) and elevated expression of inflammatory cytokines are detectable in the mediobasal hypothalamus (MBH) within the first week of HFD exposure [15–17,22], even before inflammation is detected in peripheral metabolic organs like liver and adipose tissue [16]. During HFD feeding, MBH microglia come into close apposition with POMC neurons, which is hypothesized to induce neuronal dysfunction through inflammatory signaling [21]. Indeed, microglial depletion or downregulation of microglial NF-κB signaling restores leptin sensitivity in hypothalamic neurons *in vivo* and leads to protection from HFD-associated weight gain [19,20]. Together, these data suggest that microglial activation alters obesity susceptibility through modulation of hypothalamic neuronal function [19].

In the CNS, the chemokine CX3CL1 (also known as fractalkine) is produced by neurons as both a membrane-bound full-length isoform and a soluble peptide released upon cleavage of the N-terminal domain. CX3CL1 binds a unique receptor CX3CR1 exclusively expressed by microglia, playing an important role to maintain microglial quiescence [23]. Differences in signaling of the two CX3CL1 isoforms appear to occur in neurodegenerative diseases. In mouse models of Parkinson’s disease, the soluble but not the membrane bound form provides neuroprotection [24,25]. While in a mouse model of Alzheimer’s disease, the membrane bound-form seems to have an impact in amyloid pathology [26]. Mice with reduced CX3CL1-CX3CR1 signaling (e.g. homozygous *Cx3cr1*^*gfp*^ knock-in mice) show disproportionately high microglial activation and production of inflammatory mediators, leading to increased susceptibility to CNS inflammatory pathologies such as Alzheimer’s and Parkinson’s disease [27–30]. We have previously demonstrated that MBH CX3CL1-CX3CR1 signaling decreases during HFD exposure, suggesting a potential mechanism by which HFD feeding triggers microglial activation and weight gain. Indeed, restoring CX3CL1 expression in the MBH protects from HFD-induced obesity (DIO) [20], but the mechanism linking CX3CL1 action to metabolic alterations remains unclear. Here, we provide evidence that microglia activation reduces POMC neuron excitability to promote weight gain, a mechanism reversed by CX3CL1-mediated microglial inhibition.

## 2. Materials and Methods

### 2.1 Animals

Adult male C57BL/6J and Cx3cr1-GFP [31] mice, obtained from Jackson Laboratory (JAX stock #000664, #005582), and POMC-MAPT-GFP [32], a kind gift from Dr. Tamas Horvath (Yale University), were housed in temperature-controlled rooms with 12:12 h light:dark cycle under specific-pathogen free conditions. POMC-MAPT-GFP colony was maintained as homozygotes in our animal facility and used for electrophysiological measurement of POMC neurons. All procedures were performed in accordance with NIH Guidelines for Care and Use of Animals and were approved by the Animal Care Committee at the University of Washington.

### 2.2 Reagents

Vectors containing the hybrid CMV-chicken β-actin promoter driving mRNA transcription of the soluble form of CX3CL1 (CX3CL1-s; aa 1-336) or the mutant membrane-bound form (CX3CL1-m; mutations to stop ADAM10/17 cleavage into the soluble form, R337A+R338A) with a hemagglutinin (HA)-tag appended to the C-terminus [24] (kindly provided by Dr. Kevin Nash, University of South Florida) were packaged into a AAV9 vector by the University of Washington Diabetes Research Center Viral Vector and Transgenic Mouse Core. The AAV9-GFP control virus was kindly provided by Dr. Michael W. Schwartz, University of Washington. Recombinant mouse CX3CL1 carrier-free for intracerebroventricular (i.c.v.) administration (BioLegend), and with BSA as a carrier protein to enhance protein stability for *in-vitro* studies (R&D Systems; 458-MF). Leptin was obtained from the National Hormone and Peptide Program (Dr. A. F. Parlow) and SHU9119 from Bachem.

### 2.3 Surgical procedures

#### 2.3.1 CX3CL1-s and CX3CL1-m hypothalamic overexpression

Three groups of 8- to 10-week-old male wild-type C57BL6/J mice were injected with AAV9-GFP, AAV9-CX3CL1s or AAV9-CX3CL1-m (viral titer: 1×10^12^ viral particles/ml; n=7/group). Briefly, mice from each AAV group received two consecutive stereotaxic injections (0.2 μl) bilaterally into the arcuate and ventromedial hypothalamus (anterior-posterior, −1.4 mm; lateral, ±0.5 mm; dorsal-ventral, −5.3 and −5.7 mm) using a Hamilton syringe (80030) with a 33-gauge needle at a rate of 50 nl/min (Micro4 controller) followed by a 7 min waiting period before needle removal. Mice were given 2 weeks to recover and acclimated to handling for 1 week before switching to a HFD. Body weight and food intake were recorded daily during the first week on HFD and then twice a week until the end of the study (25 days after diet switch). A leptin sensitivity study was performed in these animals (see below). The location of viral infection/protein expression was verified postmortem in all mice by immunohistochemistry (IHC) and animals that displayed marker staining outside the MBH were excluded from analysis.

#### 2.3.2 Acute SHU9119/CX3CL1 i.c.v. injections

C57BL6/J male mice underwent implantation of a lateral ventricle (LV) cannula (0.7 mm posterior to bregma; 1.3 mm lateral, and 1.3 mm below the skull surface). Mice were given 2 weeks for recovery before i.c.v. administration. The day of the experiment, four groups: 1) Saline / Saline, 2) SHU9119 / Saline, 3) Saline / CX3CL1, 4) SHU9119 / CX3CL1) of weight-match mice were established (n = 5 per group) and food was removed from at 8:00 am. At 12:00 pm mice were injected i.c.v. with saline or SHU9119 (0.5 nmol in 1μl) and 30 minutes later received a second i.c.v. injection of saline or CX3CL1 (1μg in 1μl) (See time scale in Fig. 4A). Animals were then exposed to HFD and food intake and body weight gain was measured after 24 hours.

#### 2.3.3 Chronic SHU9119 i.c.v. infusion

Two groups of 8- to 10-week-old male wild-type C57BL6/J mice were injected with AAV9-GFP or AAV9-CX3CL1-s as described above. Over the same surgical procedure, mice underwent implantation of a LV cannula (Brain Infusion Kit, ALZET) using standard stereotaxic co-ordinates (0.7 mm posterior to bregma; 1.3 mm lateral, and 2.1 mm below the skull surface). A catheter tube, filled with sterile saline, was connected to the cannula and implanted subcutaneously in the scapular region of the animal. After 10 days of recovery time from surgery, daily measurement of body weight and food intake was recorded. Two days before switching to HFD, an osmotic minipump (model 1002, Alzet) containing SHU9119 was connected to the catheter and implanted subcutaneously for 14-day i.c.v. infusion (0.2 nmol/h at 0.25 μl/h). The rate of SHU9119 infusion was based on previous studies in rodents showing that this dose effectively blocks MC4R [33,34]. Measurements of body weight and food intake were continued for 3 weeks. The location of viral infection/protein expression was verified postmortem in all mice by IHC.

### 2.4 Leptin Sensitivity

Mice (n=7/group) received daily injections of saline (200 μl/day i.p.) for 2 days followed by daily leptin (2 μg/g body weight i.p.) for 2 days with body weight and food intake measured daily [35,36].

### 2.5 Cell culture experiments

The immortalized murine microglial cell line, BV2 cells [37], was a kind gift of Dr. Suman Jayadev (University of Washington). The mouse cell line N43/5 was purchased from CELLutions Biosysems (Cedarlane Laboratory, Hornby, ON, Canada) and was derived originally from a POMC-positive fetal hypothalamic neuron generated by Dr. D. Belsham. BV2 cells were seeded at 1×10^5^ cells per well in DMEM supplemented with 10 % fetal bovine serum (FBS), 100 U/mL penicillin and 100 ug/mL streptomycin) of a 12 well plate and allowed to adhere for 16 h. Cells were serum starved for 24 h following by a 4 h incubation with recombinant CX3CL1 (50nM) or vehicle. Then, the cells were washed 3 times and incubated for another 24 h. Following incubation, medium was harvested and used as conditioned media for the proceeding cell culture experiment. Hypothalamic neuronal cells N43/5 were seeded at 1×10^5^ cells per well in a 12-well plate and treated with conditioned media from BV2 + CX3CL1 (BV2-CM-CX3CL1) or BV2 + vehicle (BV2-CM-Vehicle) for 4 h. Following incubation, cells were washed 3 times and immediately freeze at −80°C until RNA extraction. N43/5 cells were also incubated in the presence or absence of CX3CL1 (same concentration used for BV2) for 4 h to examine a direct effect of the chemokine on the neuronal cell line.

### 2.6 Brain slice preparation and electrophysiology

Ten-week-old POMC-MAPT-GFP male mice were fed chow or HFD for 4 weeks and weekly measurement of body weight and food intake was recorded. Animals were sacrificed by decapitation at the beginning of their light cycle and the fed status was confirmed by blood glucose levels taken before isofluorane anesthesia. The brains were immediately immersed in ice-cold high magnesium artificial cerebrospinal fluid (ACSF) (in mM): NaCl 125, KCl 2.5, NaHCO_3_ 26, NaH_2_PO_4_ 1.25, glucose 11, CaCl_2_ 0.5, MgCl_2_ 2 (pH 7.3, bubbled with 95% O2 and 5% CO_2_) to protect from excitotoxicity, and 300 μm acute hypothalamic slices were made using a Leica vibratome. The slices were then recovered for 30 mins at 32°C in normal ACSF containing (in mM): NaCl 125, KCl 2.5, NaHCO_3_ 26, NaH_2_PO_4_ 1.25, glucose 11, CaCl_2_ 2, MgCl_2_ 1 (pH 7.3, bubbled with 95% O2 and 5% CO2). After recovery, the slices were transferred to HEPES ACSF containing (in mM) NaCl 125, KCl 2.5, NaHCO_3_ 21, HEPES 10, Glucose 11, NaH_2_PO_4_ 1.2, CaCl_2_ 2, MgCl_2_ 2 (pH 7.4). 50nM CX3CL1 (or vehicle: PBS + 0.1% BSA) or 100 μM Minocycline (or vehicle: water) was added to the HEPES ACSF and the slices were incubated for 2 hours at 32°C. At the end of incubation period, the slices were transferred to a submerged bath in the electrophysiology rig and were perfused with HEPES ACSF with 20nM CX3CL1 or 100 μM Minocycline during recordings. POMC neurons in the arcuate nucleus were visually identified by GFP expression and subjected to whole cell patch clamp using 3-6 MΩ borosilicate glass micropipettes (Sutter Instruments, UK) filled with potassium gluconate internal containing (in mM): K-Gluconate 130, KCl 10, HEPES 10, EGTA 1, Na_2_ATP 2, Mg_2_ATP 2, Na_2_GTP 0.3. All synaptic activity was blocked using DNQX (10μM), APV (50μM), and picrotoxin (50μM) in the bath to isolate intrinsic properties of the recorded POMC neurons. Neurons were stimulated with 30 pA current step for variable durations (100 - 1000 ms) to study their evoked responses to extended depolarizing stimuli. The spontaneous excitatory postsynaptic currents (sEPSC) and spontaneous inhibitory postsynaptic currents (sIPSC) recordings were made using cesium based internal solution (in mM) Cs-Me-sulfonate 115, CsCl 20, MgCl_2_ 2.5, HEPES 10, EGTA 0.6, MgATP 4, Na_2_GTP 0.4, Na-phosphocreatine 10, with inhibitory synaptic blockers (Picrotoxin) or excitatory synaptic blockers (DNQX and APV) in the bath respectively. Electrophysiology data was acquired with a Multiclamp 700B amplifier and pClamp10 software (Molecular Devices), and sampled at 20 kHz using Digidata. Data was analyzed using Clampfit10 and custom written R scripts.

### 2.7 Tissue processing

For IHC studies, animals anesthetized using ketamine + xylazine cocktail (140mg of ketamine and 12mg of xylazine / kg of body weight) were perfused with ice-cold PBS followed by 4% paraformaldehyde (PFA) in PBS using gravitational flow. Brains were removed, post-fixed in 4% PFA overnight at 4°C, sucrose (25%) protected, cryosectioned at 30 μm in the coronal plane through the hypothalamus, and stored in freezing solution (phosphate buffer, 30 % sucrose, 30 % ethylene glycol) at −20°C for IHC staining.

For RNA extraction, animals were anesthetized using CO_2_ and rapidly decapitated. The brain was extracted and immediately frozen on dry ice. Hypothalamic blocks were dissected and stored at − 80°C until RNA extraction using RNAeasy kits (Qiagen).

For microglial cell isolation, immediately following perfusion with PBS, whole brain was dissected and dispersed into a single-cell suspension using accutase digestion at 4°C for 30 minutes. After filtration (100 μm) and centrifugation at 400g for 10 minutes, cells were resuspended in FACS buffer and transferred to a 30% Percoll solution. Cell pellet was resuspended and washed twice with FACS buffer. Microglial cells were sorted for CD11b using a FACSAria II Cell sorter (Becton-Dickinson). Microglial cells were collected directly into lysis buffer (RNeasy Micro kit, Qiagen) and stored at −80 °C until processing.

### 2.8 Real-time PCR

Total RNA was extracted using RNeasy mini kit according to manufacturers’ instructions (Qiagen) and reverse-transcribed with Multiscribe Reverse Transcriptase (Applied Biosystems). Levels of mRNA for *Pomc, Cx3cr1* and *18S* RNA (internal control) were measured by semiquantitative real-time PCR on an ABI Prism 7900 HT (Applied Biosystems).

### 2.9 Immunohistochemical Staining

Hypothalamic sections from Cx3cr1-GFP mice were incubated with either rabbit anti-Iba1 (1: 1,000; Dako), mouse monoclonal Anti-GFAP-Cy3 (1: 1,000; Sigma), mouse anti-NeuN (1: 1,000; Millipore) or chicken anti-Vimentin (1: 5,000; Abcam). After primary incubation overnight at 4ºC, all wells were processed for 1 h with appropriate fluorescent secondary antibodies (Alexa Fluor 594 donkey anti-rabbit (1: 1,500, Life Technologies); Alexa Fluor 488 donkey anti-chicken (1:500, Jackson Immuno Research); Alexa Fluor 594 goat anti-rabbit (1:500, Invitrogen); Alexa Fluor 594 goat anti-chicken (1:500, Jackson Immuno Research), Alexa Fluor 594 donkey anti-mouse (1:500, Invitrogen)). Stained sections were imaged using a cooled CCD camera attached to a DS-Ri1 epifluorescence microscope (Nikon, Japan).

### 2.10 Statistical Analyses

All results are presented as mean ± SEM. Statistical analysis using Prism (GraphPad) involved unpaired and paired two-tailed Student’s *t* tests for data with normal distribution, and Kolmogorov-Smirnov test (KS test) for data with non-normal distribution, two-way ANOVA with post-hoc Tukey’s or Sidak’s multiple comparisons testing as noted. Probability (p) values of less than 0.05 were considered statistically significant.

## 3. Results

### 3.1 Hypothalamic CX3CL1 reduces food intake and body weight and improves leptin sensitivity in HFD-fed mice

Our previous study demonstrated that DIO is associated with downregulation of CX3CL1-CX3CR1 signaling in the MBH of male mice, and that central CX3CL1 protects from diet-induced microglial inflammation and weight gain [20]. Since CX3CL1 is present in membrane-bound and soluble-cleaved forms with distinct functions in the context of neuroinflammation[24], we sought to determine whether the two isoforms have similar metabolic properties. MBH overexpression of the CX3CL1-s and CX3CL1-m isoforms was achieved by bilateral injections of AAVs containing either Cx3cl1-s, Cx3cl1-m or GFP control. Two-weeks after recovering from stereotaxic surgery, mice were placed on HFD and body weight and food intake was recorded. Reduction of weight gain and food intake over 25 days of HFD feeding was equivalent between AAV-CX3CL1-s and AAV-CX3CL1-m when compared to the AAV-GFP control (Fig. 1A-B and Supplementary Figure 1A-B). Since there was no observable difference in the metabolic effect of the two CX3CL1 isoforms, we used AAV-CX3CL1-s for the rest of our mechanistic investigations. AAV-CX3CL1-s reduced food intake during the first week of HFD (Fig. 1B), specifically between day 3 and 7 (Fig. 1C).

**Figure 1.**
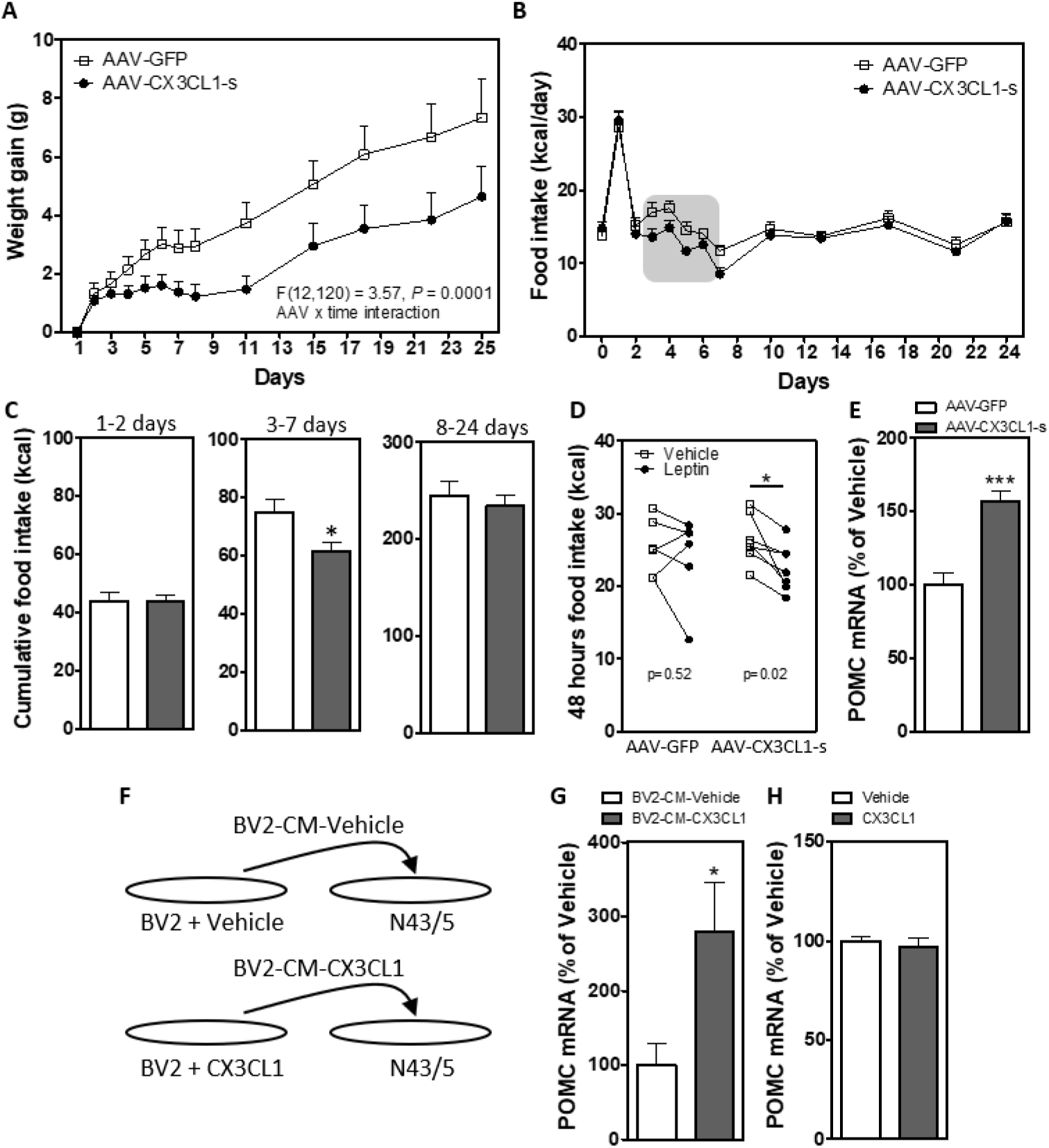
Hypothalamic overexpression of soluble CX3CL1 reduces weight gain, food intake and improves leptin sensitivity in DIO mice. **(A-B)** Body weight gain (A) and 24h food intake (B) in HFD-fed mice with hypothalamic overexpression of GFP (AAV-GFP) or the soluble isoform of CX3CL1 (AAV-CX3CL1-s) measured daily over the first week and then bi-weekly until day 25 of HFD. **(C)** Cumulative food intake over days 1-2 (left panel), 3-7 (middle panel) and 8-24 (right panel). **(D)** 48 h food intake after ip saline or leptin injections (2 μg/g/day) in AAV-GFP and AAV-CX3CL1-s mice on HFD. Data are presented as mean ± SEM of at least 6 animals per group. **(E)** Hypothalamic mRNA expression of *Pomc* in chow-fed AAV-GFP and AAV-CX3CL1-s mice. Data are expressed as a percentage of AAV-GFP control; n=6 per group. *p<0.05; ***p<0.001. **(F)** Diagram showing the cell culture experimental approach. **(G)** *Pomc* mRNA expression in hypothalamic neuronal N43/5 cells treated with CM from BV2 cells incubated with vehicle (BV2-CM-Vehicle) or CX3CL1 (BV2-CM-CX3CL1). **(H)** *Pomc* mRNA levels in N43/5 cells incubated directly with vehicle or CX3CL1. Data are expressed as a percentage of vehicle control. *p<0.05

We previously demonstrated that central CX3CL1 administration can suppress microglial activation [20]. Since HFD-induced gliosis and inflammation are associated with leptin resistance [17,19], we hypothesized that hypothalamic CX3CL1-s overexpression would increase leptin sensitivity. Indeed, once-daily injection with leptin (2 μg/g i.p.) for 2 consecutive days significantly reduced food intake in AAV-CX3CL1-s mice but not AAV-GFP controls (Fig. 1D). Consistent with its ability to improve leptin sensitivity, AAV-CX3CL1-s increased hypothalamic expression of *Pomc* in weight-matched AAV-GFP controls (Fig. 1E).

### 3.2 CX3CL1 action on microglia indirectly upregulates Pomc gene expression

CX3CR1 is the unique receptor for CX3CL1 and is expressed exclusively on microglia and a small population of perivascular macrophages within the CNS ([29,31,38,39] and Supplementary Figure 2). Thus, the effect of CX3CL1 on *Pomc* expression is most likely mediated indirectly through microglia. To address this possibility, we collected 24 hours conditioned media (CM) from murine microglia cells (BV2) previously exposed to recombinant mouse CX3CL1 (50 nM) for 4 hours. Subsequently, cultured neurons (embryonic mouse hypothalamic cell line N43/5) were exposed to CM *in vitro* for 4 hours (Fig. 1F). CM treatment increased *Pomc* mRNA expression in hypothalamic neurons (Fig. 1G) while direct exposure of N43/5 neurons to CX3CL1 had no effect (Fig. 1H). Together, these data indicate that CX3CL1 action on microglia regulates *Pomc* gene expression, suggesting close links between microglial activity and melanocortin neuron signaling.

### 3.3 Inhibition of activated microglia restores intrinsic excitability of POMC neurons in HFD-fed mice

Chronic HFD consumption is associated with hypothalamic microglial activation, close apposition of microglia to POMC neuron soma, and alterations in the electrophysiological properties of POMC neurons [4,16,17,19,21,22,40], but whether these diet-induced responses are related has not been determined. Whole-cell recordings were performed in POMC-MAPT-GFP mice, which have fluorescently-labeled POMC neurons. GFP positive POMC neurons (Fig. 2A-C) in age-matched mice exposed to chow or HFD for 4 weeks (body weight: Chow 27.65 g ± 0.74 g, N = 11; HFD 33.63 g ± 1.13 g, N = 13; p = 0.003, *t*-test) had equivalent resting membrane potential (Fig. 2D, Chow: −69.38 ± 4.3 mV, n = 11, N = 3; HFD: −70.92 ± 3.1 mV, n = 13, N = 3, p = 0.77, *t*-test). A similar proportion of POMC neurons were found to be spontaneously active in both chow and HFD fed mice (∼28%; Fig. 2E), with equivalent firing rates (Fig. 2F, Chow: 2.220 ± 0.87 Hz, n = 6, N = 5; HFD: 1.35 ± 0.58 Hz, n = 6, N = 5, p = 0.41, *t*-test). This suggests that the HFD exposure or GFP expression did not result in fundamental cellular abnormalities in the POMC neurons.

**Figure 2.**
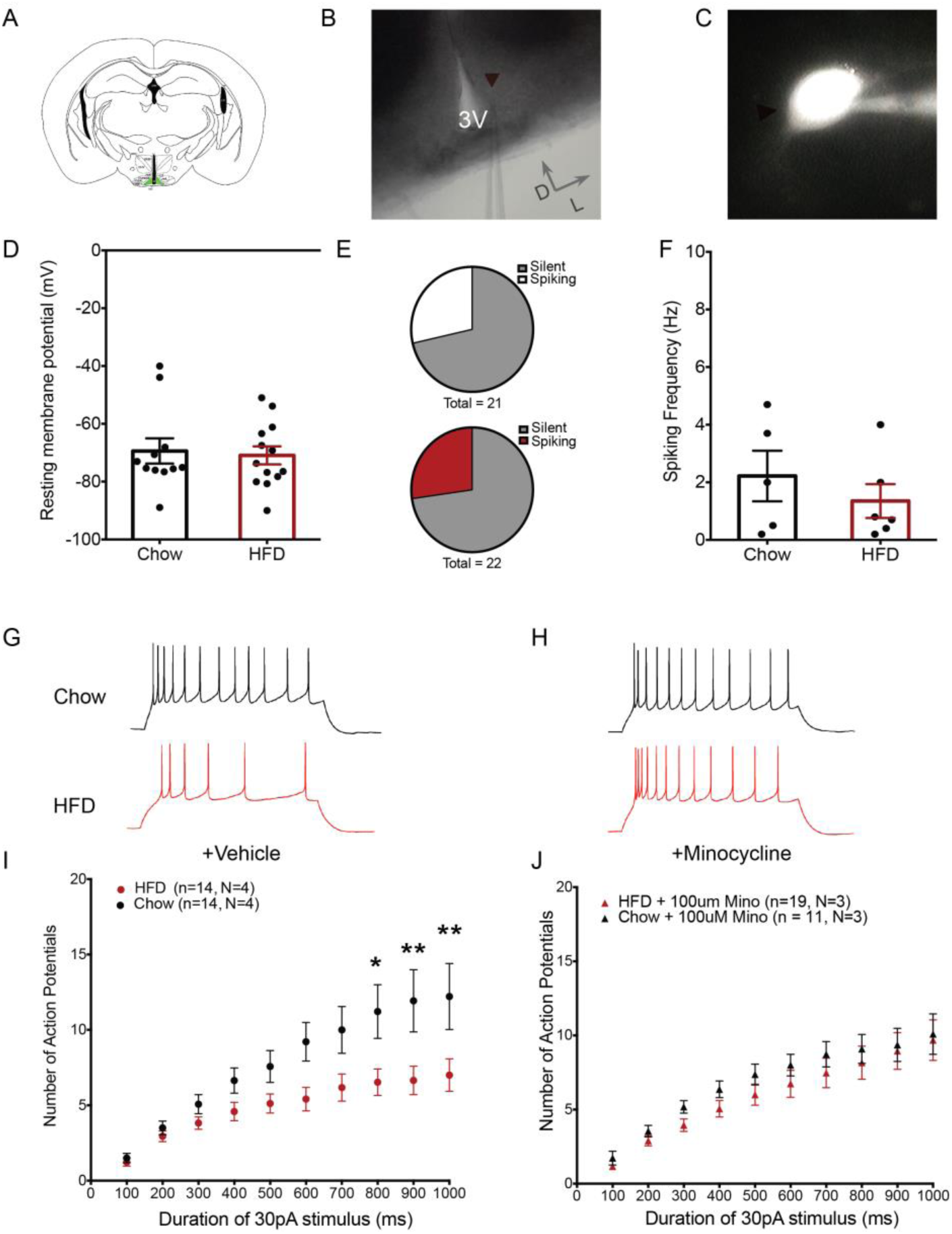
Minocycline restores intrinsic excitability of POMC neurons in HFD-fed mice. **(A)** Schematic diagram showing a coronal hypothalamic slice. The arcuate nucleus (ARC), where GFP expressing POMC neurons were visualized and recorded from, is highlighted in green. **(B)** 4X DIC image showing the ARC region and recording pipette attached to a recorded POMC neuron. **(C)** Representative fluorescent image of a GFP expressing POMC neuron with attached recording pipette filled with internal solution containing fluorophore Alexa-594. **(D)** Resting membrane potential of recorded POMC neurons (Chow: −69.38 ± 4.3 mV, n = 11, N = 3; HFD: −70.92 ± 3.1 mV, n = 13, N = 3). **(E)** Frequency of action potentials in spontaneously firing POMC neurons (Chow: 2.220 ± 0.87 Hz, n = 6, N = 5; HFD: 1.35 ± 0.58 Hz, n = 6, N = 5). **(F)** Proportion of recorded POMC neurons showing spontaneous activity (upper: Chow (6 out of 21 neurons), lower: HFD (6 out of 22 neurons)). **(G-H)** Representative traces of POMC neurons responses after stimulation by 1000 ms 30 pA current step in brain slices from chow and HFD-fed mice treated with vehicle (G) or minocycline (100μM) (H). **(I-J)** Population data of evoked responses of POMC neurons to variable duration of stimulus in brain slices from chow and HFD-fed mice incubated with vehicle (I) or minocycline (J). Lowercase indicates number of neurons and uppercase (N) number of animals. *p<0.05

Previous studies suggest that POMC neurons act as “synaptic integrators” summating electrical inputs over prolonged timescales [41]. Thus, we examined the responses of POMC neurons evoked by a minimal supra-threshold stimulus of variable duration (100 - 1000 ms, 30 pA). POMC neurons of HFD-fed mice and chow-fed mice responded with similar numbers of action potentials when stimulated by a short duration stimulus (Fig. 2G, I; 100 ms, 30 pA; Chow: 1.5 ± 0.3, n = 14, N = 3, HFD: 1.2 ± 0.26, n = 17, N = 4). However, a longer duration stimulus of the same amplitude evoked fewer action potentials in POMC neurons from HFD-fed mice compared to those from chow-fed controls (Fig. 2G, I, 1000 ms, 30 pA, Chow: 12.21 ± 2.19, n = 14, N = 3, HFD: 7 ± 1.0, n = 17, N = 4, adjusted p-value (diet) = 0.03, Two-way ANOVA, Sidak’s multiple comparisons test). This response was observed in the absence of any inhibitory or excitatory input since brain slices were incubated with blockers of glutamatergic and GABAergic postsynaptic currents, suggesting that the intrinsic excitability of the POMC neurons was significantly altered by exposure to HFD. Incubating the hypothalamic slices from HFD-fed animals in 100 μM minocycline (a tetracycline derivative that inhibits microglial activation) [42–44] for 2 hours restored the number of evoked action potentials in HFD POMC neurons to chow levels (Fig. 2 H,J, Chow: 10.09 ± 1.36, n = 11, N = 3, HFD: 9.68 ± 1.37, n = 19, N = 3, adjusted p-value (diet) = 0.40, Two-way ANOVA, Sidak’s multiple comparisons test), suggesting that microglial activation induced by HFD feeding is necessary for altering the intrinsic excitability of POMC neurons.

### 3.4 CX3CL1 restores intrinsic excitability of POMC neurons in HFD fed mice

The fact that the microglial inhibitor minocycline reversed the effect of HFD feeding on POMC neuron excitability suggested a potential mechanism explaining the metabolic benefits of CX3CL1. Like minocycline, hypothalamic CX3CL1 reduces microglial activation [20]. Additionally, we observed that CX3CL1 limited food intake, improved leptin sensitivity (Fig. 1B-D), and increased *Pomc* gene expression (Fig. 1E). Thus, we tested whether CX3CL1 treatment could restore POMC neuron excitability in HFD-fed mice. Hypothalamic slices from chow and HFD-fed POMC-MAPT-GFP mice were incubated with recombinant mouse CX3CL1 (soluble form) for 2 hours prior to recording. Similar to the effect observed with minocycline (Fig. 2K), CX3CL1 restored the number of evoked action potentials in HFD POMC neurons to chow levels (Fig. 3A-B, 1000 ms, 30 pA, Chow + CX3CL1: 11.91 ± 2.36, n = 12, N = 3; HFD + CX3CL1: 11.27 ± 1.47, n = 11, N = 3, HFD + Veh: 6.70 ± 1.43, n = 10, N = 3, adjusted p-value (diet+intervention) = 0.08, Two-way ANOVA, Sidak’s multiple comparisons test).

**Figure 3.**
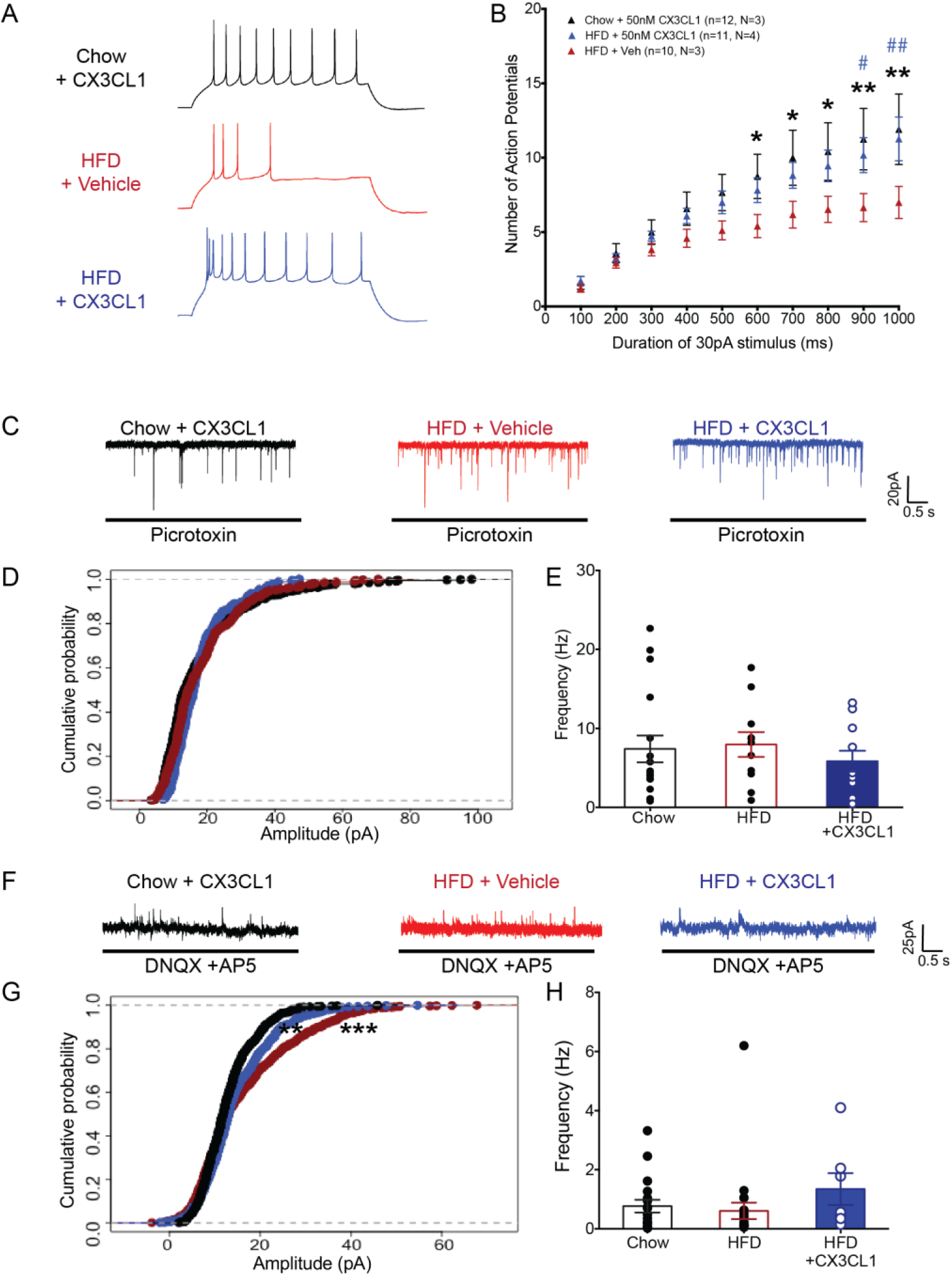
CX3CL1 restores intrinsic excitability of POMC neurons in DIO mice. **(A)** Representative traces of POMC neurons after stimulation by 1000ms 30pA current step (top: stimulus step; bottom: responses: black: Chow + CX3CL1, red: HFD + Veh, blue: HFD + CX3CL1(* denotes p values for Chow+CX3CL1 and HFD+veh comparison, # denotes p values for HFD+veh and HFD+CX3CL1 comparison)). **(B)** Population data of evoked responses of POMC neurons to variable duration of stimulus after CX3CL1 treatment in brain slices from chow and HFD-fed mice. **(C-E)** Representative traces (C), cumulative probability plot of amplitude (D), and frequency plot (E) of sEPSCs for POMC neurons recorded in chow and HFD brain slices incubated for 2 hours with vehicle or CX3CL1 (Frequency of sEPSCs: Chow + CX3CL1: 7.41 ± 1.69 Hz, n = 17, N = 3; HFD + Veh: 7.96 ± 1.56 Hz, n = 11, N = 3; HFD + CX3CL1: 5.84 ± 1.32 Hz, n = 11, N = 3). **(F-H)** Representative traces (F), cumulative probability plot of amplitude (G), and frequency plot (H) of sIPSCs for POMC neurons recorded in chow and HFD brain slices incubated for 2 hours with vehicle or CX3CL1 (Frequency of sIPSCs: Chow + Veh: 0.76 ± 0.21 Hz, n = 18, N = 4; HFD + Veh: 0.6 ± 0.27 Hz, n = 22, N= 4; HFD + CX3CL1: 1.3 ± 0.5, n = 7, N = 2). Lowercase (n) indicates number of neurons and uppercase (N) number of animals. *p<0.05; **p<0.01; ***p<0.001

We also assessed the effect of CX3CL1 treatment on synaptic inputs to POMC neurons. Spontaneous excitatory postsynaptic currents (sEPSCs) received by POMC neurons remained unchanged in HFD as well as CX3CL1-treated brain slices compared to chow (Fig 3C-E, Chow: 7.41 ± 1.69 Hz, n = 17, N = 3; HFD: 7.98 ± 1.56 Hz, n = 11, N = 3; HFD + CX3CL1: 5.84 ± 1.32 Hz, n = 11, N = 3). Likewise, there was no difference in the frequency of spontaneous inhibitory postsynaptic currents (sIPSCs) in POMC neurons from HFD vs chow-fed mice (Fig. 3F, H, Chow: 0.76 ± 0.21 Hz, n = 18, N = 4; HFD: 0.6 ± 0.27 Hz, n = 22, N = 4; HFD + CX3CL1: 1.3 ± 0.5, n = 7, N = 2, p-value = 0.66, t-test). Interestingly, HFD exposure increased the cumulative probability of larger amplitude sIPSCs in POMC neurons suggesting increased inhibition on POMC neurons due to exposure to HFD (Fig 3G, red trace, chow vs HFD: p value = 6.37e-12, KS test). CX3CL1 application was able to partially restore the inhibition to chow levels (Fig. 3G, blue trace, HFD+veh vs HFD+CX3CL1: p value = 0.0015, KS test). Taken together, these results support the hypothesis that microglial CX3CL1- CX3CR1 signaling reverses the impairment of hypothalamic POMC neuronal activity induced by HFD feeding.

### 3.5 The anti-obesity effect of central CX3CL1 requires intact melanocortin signaling in vivo

The degree of POMC neuron activity generally correlates with production of α-MSH, a neuropeptide derived from POMC that reduces food intake by acting on MC3R and MC4R [45,46]. Thus, the ability of CX3CL1 to restore POMC neuron excitability suggested a potential effect to increase melanocortin signaling during DIO (also indicated by the increase in *Pomc* gene expression (Fig. 1E)). We investigated whether pharmacological blockade of the central melanocortin signaling could impede the ability of CX3CL1-s to reduce weight gain and food intake in HFD-fed mice. A single i.c.v. administration of recombinant mouse CX3CL1 peptide (soluble form) tended to reduce 24-hour HFD intake (Fig. 4B, Sal/Sal vs Sal/CX3CL1) and significantly limited weight gain (Fig. 4C, Sal/Sal vs Sal/CX3CL1). Pre-treatment with a dose of the MC3R/MC4R antagonist SHU9119 (0.1 nmol, i.c.v.) that by itself had no effect on food intake and body weight (Fig. 4B-C, Shu/Sal vs Sal/Sal) blocked the metabolic benefits of acute CX3CL1 treatment (Fig. 4B-C, Sal/CX3CL1 vs Shu/CX3CL1). To assess the involvement of melanocortin system activation in DIO prevention by CX3CL1, s.c. osmotic minipumps containing SHU9119 (0.2 nmol/h) were placed in lateral ventricle-cannulated mice with prior AAV-CX3CL1-s or AAV-GFP injection into the MBH (Fig. 4D). While animals with hypothalamic overexpression of CX3CL1-s have reduced food intake and body weight during HFD feeding (Fig. 1A-C), concurrent SHU9119 infusion abolished these benefits (Fig. 4E-F). Collectively, these data support a requirement for intact melanocortin signaling for the anti-obesity effect of hypothalamic CX3CL1.

**Figure 4.**
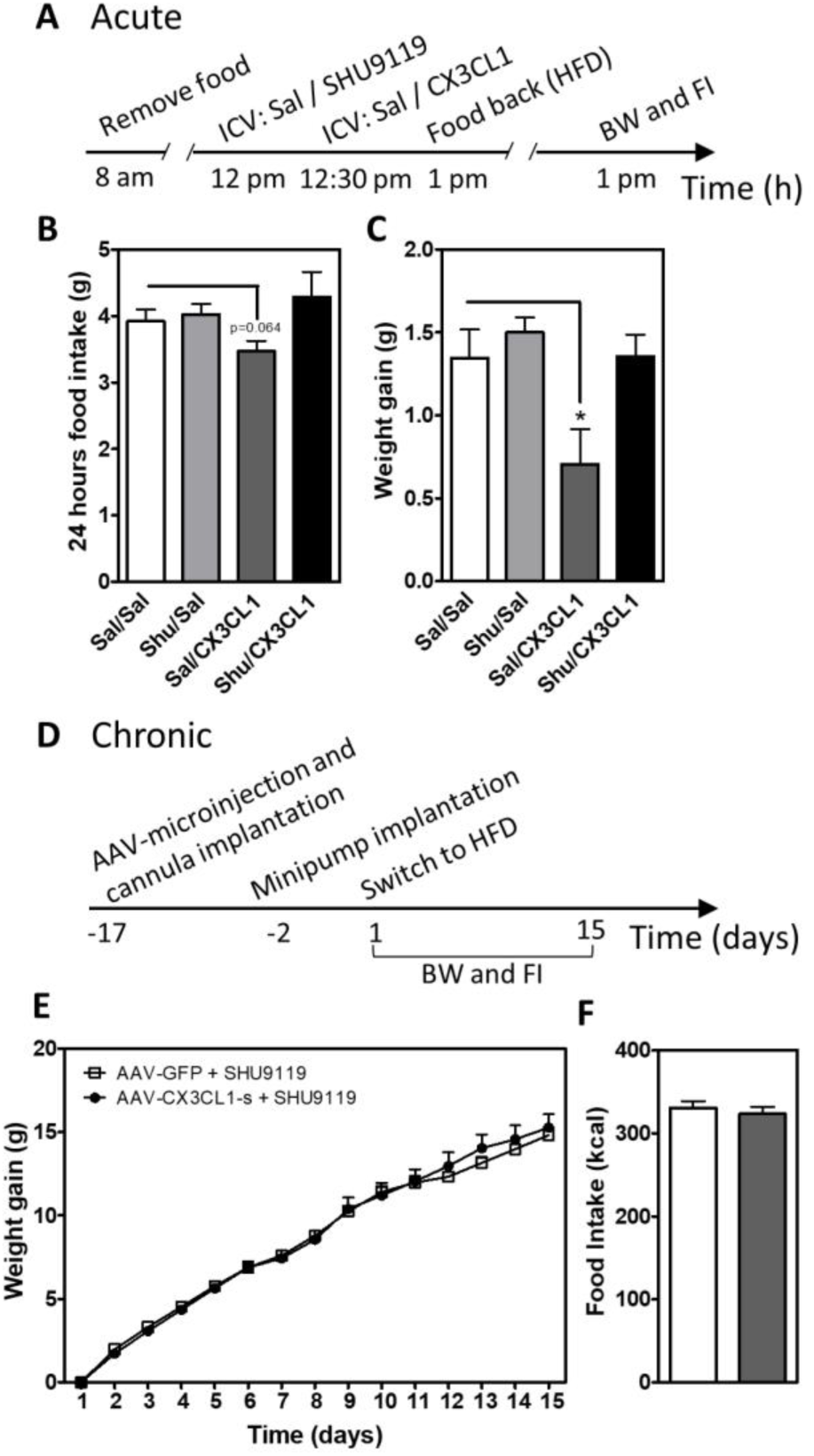
Central CX3CL1 does not reduce weight gain and food intake in mice treated with MC3R/MC4R antagonist. **(A)** Time scale of the experimental approach used for acute i.c.v. administration of SHU9119 and CX3CL1. **(B-C)** Average of cumulative 24 h food intake (A) and body weight gain (B) in mice treated with saline (Sal) or SHU9119 i.c.v. and 30 minutes later with Sal or CX3CL1 i.c.v. Data are presented as mean ± SEM of at least 5 animals per group. *p<0.05. **(D)** Time scale for chronic i.c.v. administration of SHU9119 in mice previously subjected to hypothalamic viral microinjection of AAV-GFP and AAV-CX3CL1. **(E-F)** Body weight gain (E) and total food intake (F) measured in mice with hypothalamic overexpression of CX3CL1-s or GFP control that received SHU9119 i.c.v. infusion over 2 weeks. Data are presented as mean ± SEM of 10 animals per group. *p<0.05.

## 4. Discussion

Neuronal output dictates behavior including food intake and metabolic rate, yet diet-induced activation of glial cells can promote susceptibility to hyperphagia and obesity [3,16,18,19,47,48]. Therefore, glial cells must influence energy balance through action on neighboring neurons, but how this occurs remains unclear. Here, we provide evidence that neuronal-microglial crosstalk via CX3CL1-CX3CR1 signaling maintains melanocortin system activity during HFD feeding to protect from DIO.

CX3CL1-CX3CR1 signaling is implicated in maintaining microglia in their basal surveillance mode as well as modulating neuronal function and survival in the adult brain [49]. CX3CR1 is a G_i_ protein-coupled receptor that regulates apoptotic, proliferative, transcriptional, and migratory functions in microglia [50]. Perturbation of CX3CL1-CX3CR1 signaling is associated with increased neuroinflammation and neuronal cell death in animal models of neurodegenerative disease [24,25,29,51,52]. In the context of DIO, where microglial inflammation in the MBH has been shown to be required for body weight gain upon HFD exposure [19,47,48], we have previously shown that CX3CL1 is reduced in the hypothalamus of male mice, and that central administration of CX3CL1 prevented diet-induced weight gain and microglial activation [20]. However, the mechanisms by which this chemokine provides protection from weight gain were not addressed. Though previous studies have demonstrated different signaling functions for the membrane-bound and soluble isoforms of CX3CL1 in neurodegenerative conditions [24], we found both equally effective in preventing weight gain. This occurred despite the presence of endogenous CX3CL1 (at low levels due to HFD feeding), suggesting that CX3CL1 overexpression could have pharmacological benefits for obesity treatment.

Mice overexpressing CX3CL1 in the MBH displayed a significant reduction in food intake and increase the anorexigenic response to leptin, suggesting that microglial CX3CR1 signaling may affect neurons involved in feeding regulation. Indeed, we found that CX3CL1 increases the expression of the neuropeptide POMC *in vivo*, and showed this effect to be mediated through action on microglia rather than directly on neurons. In support of this cell-cell interaction model, a previous study demonstrated an increase in microglia-POMC neuron cell contact when mice were exposed to a high-fat, high-sucrose diet [21]. The specificity of CX3CL1 action on microglia is further supported by our immunohistological analysis of Cx3cr1-Gfp mice confirming that CX3CR1 is not expressed in neurons, astrocytes, or tanycytes. Similarly, single-cell RNAseq analysis of mouse hypothalamic tissue has shown that CX3CR1 is not found in any neuronal populations but is exclusively expressed in microglia [53] as well as subsets of peripheral monocytes and macrophages [31]. Since circulating myeloid cells infiltrate the MBH and contribute to hypothalamic microgliosis in HFD-fed mice [19], the contribution of CX3CL1 action on other subsets of myeloid cells to its anti-obesity effects has not been excluded.

CX3CL1 acts as an anti-inflammatory molecule by downregulating microglial IL-1β, TNF-α and, IL-6 production [20,54,55]. We recently showed that central CX3CL1 administration to HFD-fed mice reduces the expression of TNFα in hypothalamic microglia [20]. There is accumulating evidence that TNFα is an important pro-obesogenic factor. Knockdown of TNFα signaling in MBH neurons of DIO mice reduces food intake and body weight gain while lowering mitochondrial activity and cellular stress in POMC neurons [21]. Thus, we speculate that hypothalamic CX3CL1 may restore POMC neuron excitability and limit weight gain in DIO mice by reducing TNFα signaling. However, the proinflammatory agent LPS also triggers microglial activation but increases POMC electrophysiological activity [44]. Thus, an important priority for future research is to define the molecular characteristics of HFD-induced microglial activation beyond just inflammatory pathways. In this vein, an alternative to the reduced TNF signaling model posits that CX3CL1 slows mitochondrial metabolism and reactive oxygen species production in microglia, thereby reducing damage to neighboring neurons such as closely apposed POMC cells [21]. In support of this hypothesis, microglia-specific deletion of UCP2, a critical protein in mitochondrial function, prevents HFD-induced obesity and increases POMC neuronal activity in the hypothalamus [47].

Hypothalamic inflammation and gliosis are proposed to mediate the synaptic input rearrangement, electrophysiological alterations, and neuronal loss observed in POMC neurons during HFD feeding [3,16,21]. Consistent with previous studies, we found that mice exposed to HFD have reduced POMC neuron activity due to alterations in intrinsic excitability rather than changes in synaptic inputs [4]. Minocycline treatment restored neuronal excitability, suggesting that microglial activation mediates the effect of HFD to inhibit POMC neurons. However, these data should be interpreted with caution as minocycline has been reported to affect other cell types [56]. Nevertheless, restoration of intrinsic excitability of POMC neurons in HFD fed mice by CX3CL1 support a role for microglia in POMC neuron activity during DIO. Though CX3CL1-CX3CR1 signaling is known to modulate neuronal activity in other contexts, the mechanisms involved are not completely understood [57]. One possibility is suggested by a recent study demonstrating that reduced mitochondrial Ca^2+^ uptake during HFD feeding contributes to impaired POMC neuron excitability [4]. Activation of CX3CR1 signaling may alter the microglial secretome to restore POMC neuron activity through improved mitochondrial Ca^+2^ dynamics.

In our findings, CX3CL1 treatment of HFD-fed mice improved the anorexigenic response to leptin, restored POMC neuron excitability, and increased *Pomc* expression, all actions that would be predicted to raise the level of the melanocortin agonist, α-MSH, which acts at the MC4R to reduce food intake [58]. Indeed, both acute and chronic administration of SHU9119, an MC3/4 receptor antagonist, eliminated the protective effect of CX3CL1 on diet-induced body weight gain and food intake. Though the acute effect of icv-injected CX3CL1 may involve extrahypothalamic regions including brain areas controlling food reward and motivation (important factors in the first 24 h of HFD feeding [59]), the administration of a subthreshold dose of SHU9119 still blocked the acute benefits of CX3CL1. Similarly, chronic inhibition of the melanocortin system by central infusion of SHU9119 eliminated the anti-obesity effects of overexpressing CX3CL1 in the MBH. Together, these data support the model that CX3CL1 acts through microglial inhibition to maintain melanocortin system activity during DIO.

In summary, we demonstrate that both soluble and membrane-bound CX3CL1 potently reduce weight gain when overexpressed in the MBH. CX3CL1 not only increases leptin sensitivity in DIO mice, but also reverses HFD-induced reductions in POMC neuron excitability. Moreover, the anti-obesity effect of CX3CL1 is blocked in the presence of a melanocortin receptor 3/4 antagonist. Together, these findings highlight a critical role in obesity pathogenesis for POMC neuron dysfunction triggered by microglial activation and demonstrate the therapeutic benefit of microglial inhibition using CX3CL1.

## ACKNOWLEDGEMENTS

The authors acknowledge the technical assistance provided by Lee Shaffer at the University of Washington and many discussions with members of the M.W. Schwartz laboratory. The authors are also grateful to Dr. David Morgan (University of South Florida) for providing constructs containing mutant forms of CX3CL1 (secreted and membrane chemokine domain-only forms), and Dr. Tamas Horvath (Yale University) for the *Pomc-MAPT-GFP* mice. This work was supported by the Scientist Development Award from the American Heart Association (AHA; 16SDG27010018), and two Pilot and Feasibility Awards by the Diabetes Research Center (DRC; P30 DK017047) and University of Washington Medicine Diabetes Institute (UWMDI), and UW Royalty Research Fund Award to M.D.D., by ADA Pathway to Stop Diabetes Grant 1-14-ACE-51 and R01DK119754 to J.P.T. In addition, services and support were provided by the Nutrition Obesity Research Center (DK035816) and Diabetes Research Center (DK017047) at the University of Washington.

## Author Contributions

J.B., M.D.D., A.B., and J.P.T. conceived the project and designed the research plan. J.B., M.D.D., R.D.F., J.D.D, and A.W.J performed the experiments. J.B., M.D.D., A.B., and J.P.T analyzed the results and wrote the manuscript. All the authors edited and approved the final manuscript.

## Guarantor statement

Dr. Joshua P. Thaler is the guarantor of this work and, as such, had full access to all aspects of the study and takes responsibility for the data and the accuracy of the data analysis.

## Figure Legends

**Supplementary Figure 1.**
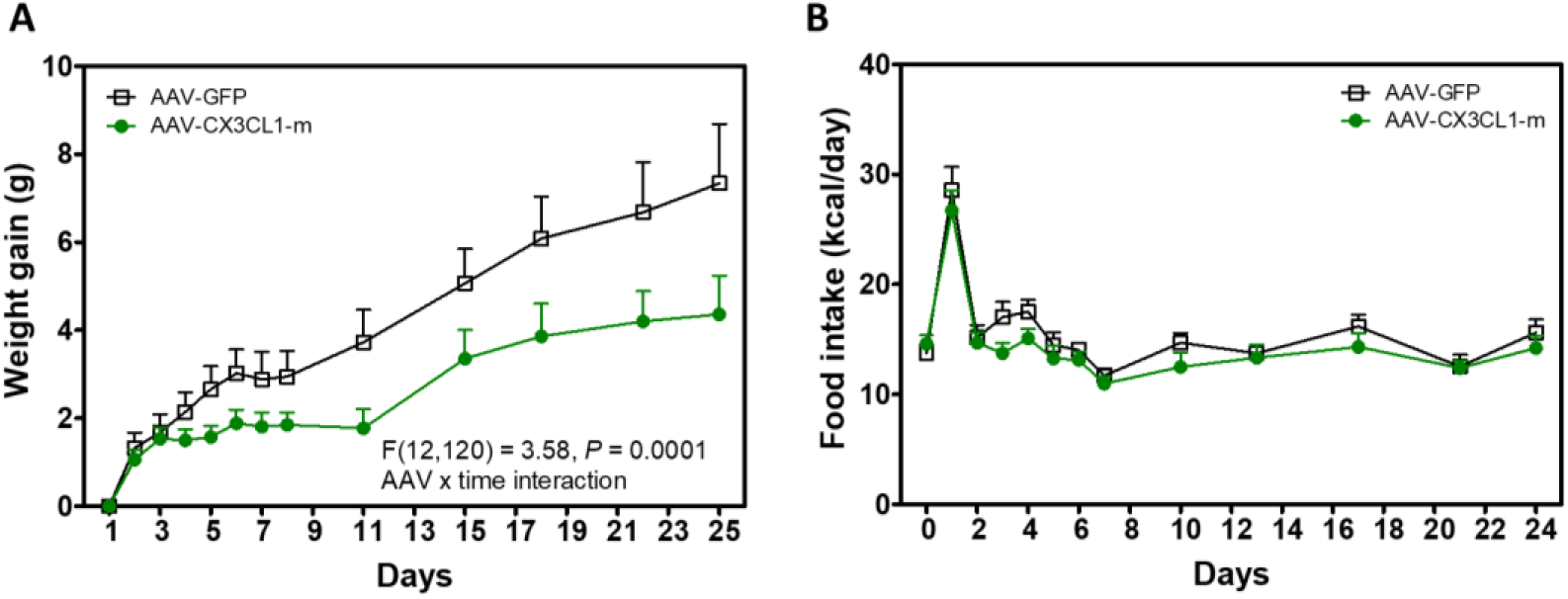
Hypothalamic overexpression of the membrane-bound isoform of CX3CL1 limits weight gain and reduces food intake in DIO mice. (A-B) Body weight gain (A) and daily food intake (B) in HFD-fed mice with hypothalamic overexpression of GFP (AAV-GFP) or the membrane-bound isoform of CX3CL1 (AAV-CX3CL1-m) measured daily over the first week and then bi-weekly until day 25 of HFD. Data are presented as mean ± SEM of at least 6 animals per group.

**Supplementary Figure 2.**
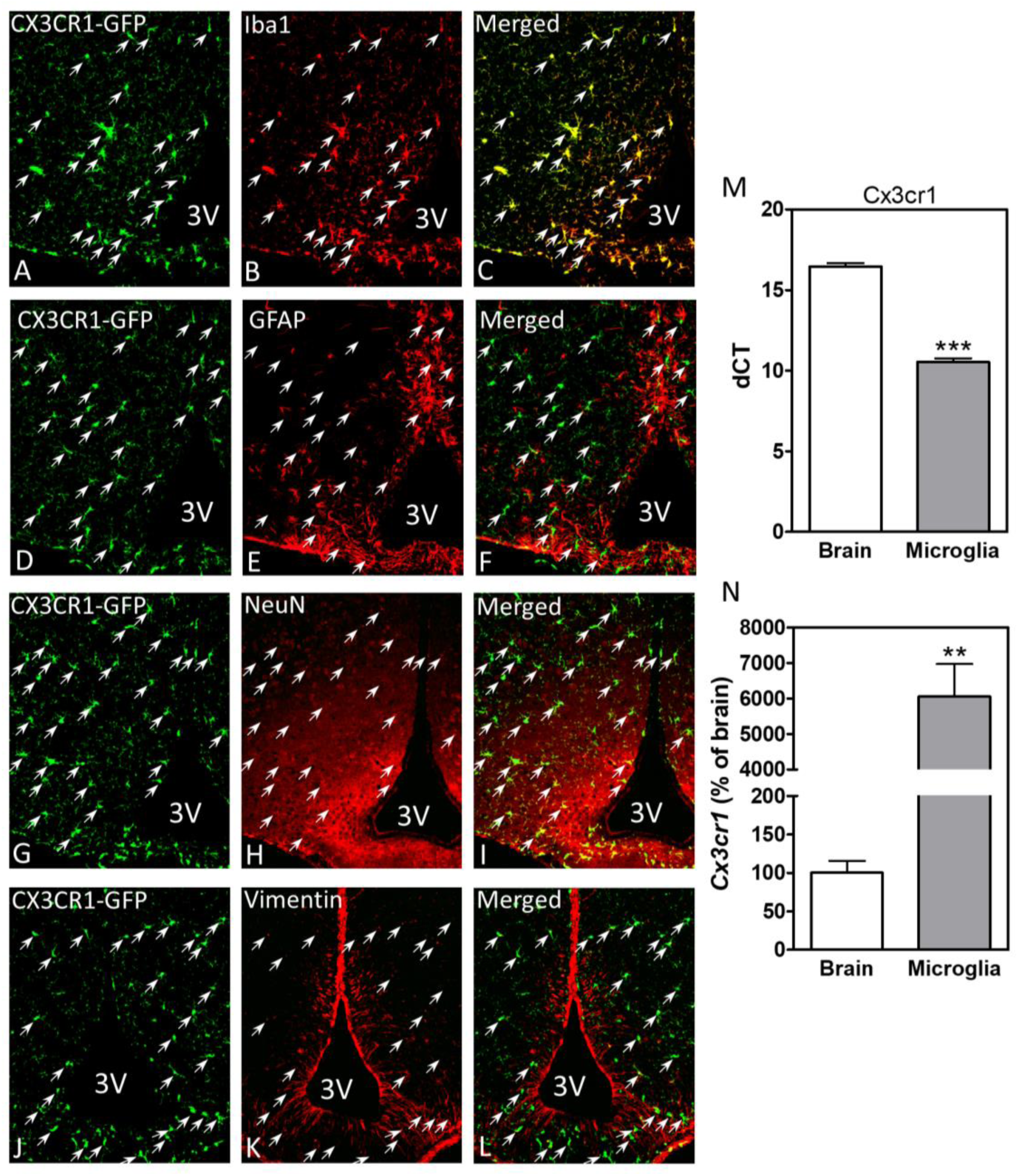
CX3CR1 is expressed in microglia, but not astrocytes, neurons or tanycytes in the MBH. Representative images showing CX3CR1-GFP (A, D, G, J) and Iba1 (B), GFAP (E), NeuN (H), Vimentin (K) immunoreactivity in the MBH. (C) All CX3CR1-GFP positive cells are also positive for the microglial marker Iba1. There is no colocalization between CX3CR1-GFP and the astrocytic marker GFAP (F), the neuronal marker NeuN (I) or the tanycyte marker vimentin (L). Delta Ct values (M) and relative expression (N) of *Cx3cr1* expressed in whole brain and sorted brain microglia. **p<0.01; ***p<0.001

